# Modeling the variability of the electromotor command system of pulse electric fish

**DOI:** 10.1101/2020.06.09.142083

**Authors:** Ángel Lareo, Pablo Varona, Francisco B. Rodríguez

**Affiliations:** Grupo de Neurocomputación Biológica, Departamento de Ingeniería Informática, Escuela Politécnica Superior, Universidad Autónoma de Madrid, Spain

## Abstract

Mormyrids, a family of weakly electric fish, use electric pulses for communication and for extracting information from the environment (active electroreception). The electromotor system controls the timing of pulse generation. Ethological studies have described several sequences of pulse intervals (SPIs) which are related to distinct behaviors (e.g. mating or exploratory behaviors). Accelerations, scallops, rasps, and cessations are four SPI patterns reported in these fish, each showing characteristic temporal structures and large variability. This paper presents a computational model of the electromotor command circuit that reproduces SPI patterns as a function of the inputs to the model while keeping the same internal network configuration. The topology of the model is based on a simplified representation of the network with four neuron clusters (nuclei). An initial *ad hoc* tuned configuration (S-T) was built to reproduce nucleus characteristics and network topology as described by detailed morphological and electrophysiological studies. Then, a genetic algorithm (GA) was developed and applied to automatically tune the synaptic parameters of the model connectivity. Two different configurations obtained from the GA are presented here: one optimized to a set of synthetic examples of SPI patterns (S-GA) and another configuration adjusted to patterns recorded from freely-behaving *Gnathonemus Petersii* specimens (R-GA). Robustness analyses to input variability were performed to discard overfitting and assess validity. Results show that the set of SPI patterns are consistently reproduced, both for synthetic data and for recorded data. This model can be used as a tool to test novel hypotheses regarding temporal structure in electrogeneration.

## 1 Introduction

Pulse mormyriforms, a group of weakly electric fish, produce electric pulses with high temporal precision. These fish have the ability of polarize their body in fast voltage transients whose deflection in the fish surroundings is detected by the fish using a specialized electric organ [5]. The electric organ discharges (EODs) occur as a result of the synchronous activation of modified muscle or nerve cells named electrocytes. This ability, known as active electroreception is a well-suited sensory modality to study information processing in a living neural system as the signal from a freely-behaving fish can be monitored during long time periods.

Information is encoded in the fish signal using a multiplexed temporal coding [2]. The pulse shape, with a mean duration of*∼*1 ms, is stereotyped, although there are variations among species [17], sex [3] or relative dominance [12]. The interval between EODs, known as the inter-pulse interval (IPI), is much larger and variable than the duration of the EOD. At rest, IPIs are around 100 to 300 ms, but they fluctuate from less than 10 to more than 400 ms [32]. IPIs and sequences of pulse intervals (SPIs) are also relevant to information processing in these animals, as complex higher-level information can be encoded using this kind of temporal coding [2, 17]. For instance, IPIs decrease when the fish is actively probing their environment [33]. The timing flexibility of IPIs in this system gives rise to an interesting set of SPI patterns with behavioral relevance, as we will discuss below.

The neural system responsible for controlling the timing of the EODs is the electromotor system, located at the central nervous system of the fish [6]. A neural ensemble known as the command nucleus (CN) initiates the EOD. Action potentials in CN are correlated with EODs (i.e., each action potential in CN leads to an EOD). Nevertheless, CN is not a pacemaker but an integrator system. It mainly receives synaptic input from the mesencephalic precommand nucleus (PCN) and the adjacent thalamic dorsal posterior nucleus (DP) (Fig. 1). Nuclei DP and PCN receive projections from multiple sources, but they are both inhibited by the ventroposterior nucleus (VPd). This inhibition is mediated by the activation of VPd through a feedback mechanism, the corollary discharge pathway (see *E*_*P CN*_ in Fig. 1). Inhibition feedback is a mechanism for avoiding responses to the fish own EOD and seems to regulate the resting electromotor rhythm [10]. This is also emphasized by the fact that the IPI of DP/PCN nuclei last, at least, as much as VPd bursts [9].

**Figure 1.**
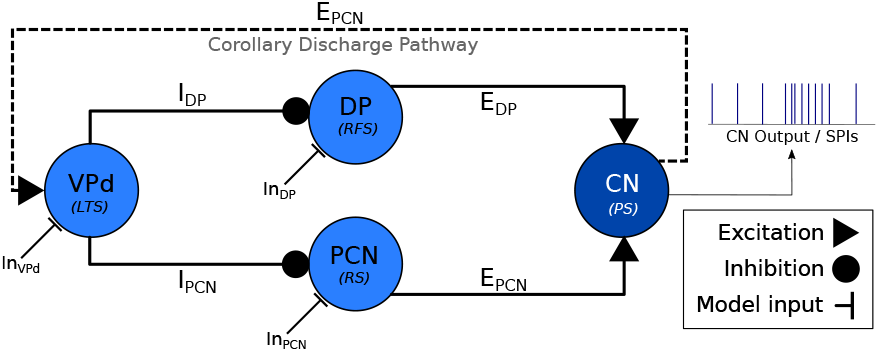
Abridged schematic of the electromotor command network based on [4, 9] and used for developing the computational model discussed in this paper. An EOD occurs after each action potential in CN [16], so CN activations represent the output of the model. CN receives excitatory projections from DP (E_DP_) and PCN (E_PCN_). Inhibitory afferents are driven to DP and PCN through VPd (I_DP_ and I_PCN_ respectively) triggered by the corollary discharge pathway (E_CDP_), which makes VPd to fire a burst of action potentials right after the production of an EOD [8]. This inhibition feedback seems to regulate the rythm of EOD output [34]. Current inputs to the electromotor model (In_VPd_, In_DP_, In_PCN_) can be tuned to simulate different behavioral conditions.

We have developed a model of the electromotor command system of pulse mormyrids capable of reproducing the variability of temporal firing patterns shown by these fish as a function of the input while sustaining the same network architecture. The model topology and nucleus dynamics are based in the results from previous physiological studies of the pulse mormyrid electromotor system [6].

Samples of these patterns were obtained for this work from experimental data recordings of living *Gnathonemus Petersii* specimens [14, 15, 21] for the first time. These types of patterns have been previously characterized in another species of the *Mormyridae* family [11].

An automated method based on genetic algorithms (GA), with the development of a multiobjective fitness function specifically adapted to this system, was applied for synaptic parameter setting. Patterns recorded from *Gnathonemus petersii* and synthetic samples generated from previously described behavioral patterns in pulse mormyrids were used to fit the model parameters. Finally, the robustness of the model was tested under systematic variations of network inputs to assess validity and discard overfitting to these inputs.

## 2 Computational model of the electromotor command network

Figure 1 illustrates the model of the electromotor command network described in the following subsections. We also describe below the genetic approach used to fit the connectivity parameters that give rise to the generation of distinct activity patterns as a function of the stimuli.

### 2.1 Characteristic sequences of pulse interval (SPIs): Synthetic and experimental data

Pulse mormyrids generate a wide variety of electrical activity patterns using different sequences of pulse intervals (SPIs). Resting IPIs range from *∼*100-300 msec. Previous studies have described several stereotyped SPIs related to distinctive social behaviors [11]. Accelerations, scallops, rasps and cessations are four relevant SPIs that have been described [6, 11, 22].

Accelerations are sustained interval shortenings to a series of regular smaller IPIs with variable duration. Accelerations (see Fig. 2) are related to the activation of the neural ensemble known as the adjacent thalamic dorsal posterior nucleus (DP, see Fig. 1) [10]. According to [20], this kind of SPI is related with aggressive behaviors.

**Figure 2.**
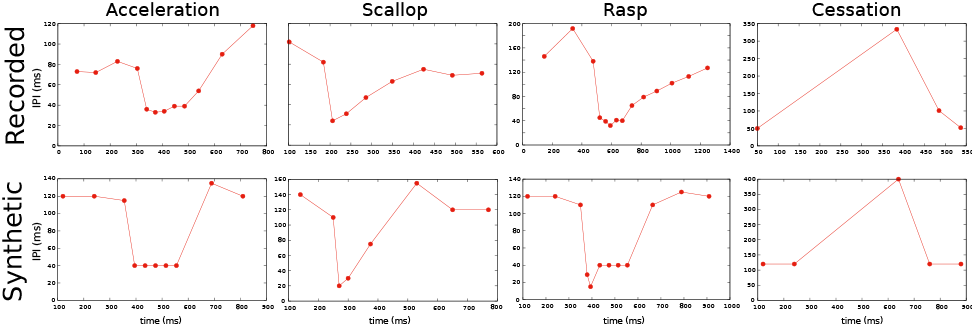
Characteristic SPI patterns recorded from freely-behaving *Gnathonemus Petersii* specimens (top) and examples of corresponding synthetic SPIs to evolve and validate the electromotor command network model (bottom). Signals were recorded using the platform and the methodology explained in the Experimental Methods section. Synthetic SPIs were constructed preserving the characteristic temporal structure of reported experimental observations [6, 10, 26].

Scallops (see Fig. 2) are sudden drops to very short IPIs followed by an almost immediate recovery to regular resting IPIs. Contrary to what occurs in accelerations, the emmission of shorter IPIs is not sustained. An scallop pattern in CN takes place after the activation of the mesencephalic precommand nucleus (PCN, see Fig. 1), and this kind of firing pattern may serve as an advertisement signal [29].

Rasps (see Fig. 2) are a type of IPI pattern that has an initial sudden decrease to very short IPIs, similar to the one observed in scallops, followed by a step increase of IPIs duration for a sustained series of IPIs, like a long tail of short regular IPIs similar to the ones observed in the acceleration pattern [18]. Rasps likely relies on activation of both DP and PCN [6] (see Fig. 1). This SPI is evoked during male courtship behavior [11].

Finally, cessations (see Fig. 2) correspond to activity dropping in the EOD generation for time periods up to one second. A cessation is evoked by the activation of the ventroposterior nucleus (VPd, see Fig. 1). Submissive behavior has been associated with this SPI [25, 26]. We will use these four representative patterns to validate our our electromotor command network modeling approach.

### 2.2 Nuclei model

Four different neuron ensembles in the electromotor command chain were modeled: The medullary command nucleus (CN), the mesencephalic precommand nucleus (PCN), the adjacent thalamic dorsal posterior nucleus (DP) and the dorsal region of the ventroposterior nucleus (VPd). Each nucleus was simulated using the neuron model developed by Izhikevich (2003) [19]. This model combines biologically plausibility and computational performance characteristic of integrate-and-fire neuron modeling approaches [23]. It is based in a two-dimensional system of ordinary differential equations:

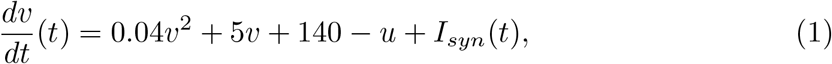

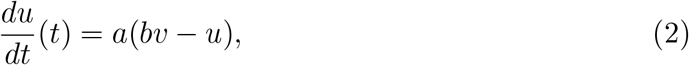

with an auxiliary after-spike resetting:

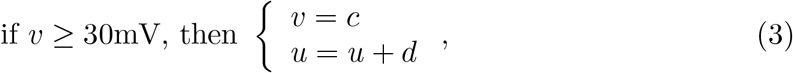

where *v* represents the neuron’s membrane voltage and *u* represents the combined action of ionic current dynamics. The parameters *a, b, c* and *d* set the working regime of the neuron model. *I*_*n*_ is the model external input.

A wide range of neuron dynamics, and in particular firing temporal structures, can be reproduced by selecting different values of the parameters *a, b, c* and *d*, as shown in [19].

The parameters of the neuron model were first adjusted to reproduce dynamics of the nuclei described by previous neurophysiological studies of the electromotor command network [9]. DP model parameters were adjusted to show regular frequency spiking, whereas PCN parameters were chosen to reproduce decreasing frequency spiking. These dynamics were selected because they are observed in the corresponding pattern evoked by each nucleus. According to [9], units from DP/PCN nuclei showed wide variations in baseline frequency, from sporadical firing units to units with high spiking rates, so no baseline firing frequency was preselected for the DP/PCN models. Nevertheless, as it occurs in the living network, a bimodal structure arose from the IPIs intervals before and after CN action potential in both DP/PCN, with larger intervals ocurring after the activation of CN. VPd nucleus model fires high-frequency sequences of action potentials with a noticeable spike frequency adaptation, which is in accordance to physiological records showing bursts of action potentials from VPd during each IPI. This behavior was modelled using a *low-threshold spiking* firing regime, which is usually displayed by inhibitory neurons (initial synaptic parameters were also adjusted to reproduce this behavior). Also, there was considerable variation in the timing of the first spike of the burst relative to CN, intraburst firing rate, burst duration, and number of spikes per burst from VPd units, so none of these characteristics were *a priori* selected. Finally, CN model was configured to a *phasic spiking* firing regime as it mainly integrates inputs from DP and PCN [7].

The parameter values for modelling each nucleus are listed in B, Table 2 depicts the activity of each model nucleus (isolated from the network) in response to a step current input.

### 2.3 Synapses model

We reproduced neural projections using a model of chemical synapses as chemical inter-nuclei communication takes place in the real electromotor command network. In these synapses, when the pre-synaptic target generates an action potential a certain amount of neurotransmitters are released and bind to the postsynaptic receptors. The mathematical description used to model this behavior is based in a description of the receptor bindings [13]. These equations define a simple method for computing synaptic currents with low computational cost. The ratio of bound chemical receptors in the post-synaptic target (*r*) during a pulse (*t*_*f*_ *< t < t*_*r*_) and after the pulse (*t*_*r*_ *< t*) was calculated as follows:

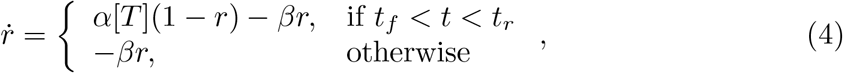

where *α* and *β* are the forward and backward rate constants for transmitter binding and [T] is the neurotransmitter concentration.

The beginning of a pulse (*t*_*f*_) was detected when presynaptic neuron’s membrane potential crossed a given *threshold*. Time between *t*_*f*_ and *t*_*r*_ was defined as the maximum release time (*t*_*max*_). Both *threshold* and *t*_*max*_ were tuned as a parameter of the synapsis.

From the ratio of bound receptors is given by equation 4, the current received by the post-synaptic target, *I*_*syn*_, at any time *t* was then calculated as follows:

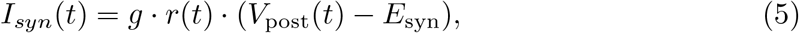

where *g* is the synaptic conductance, *V*_post_(*t*) is the post-synaptic potential at time *t*, and *E*_syn_ is the synaptic reversal potential at the same time.

The topology of the model was set up using a standard configuration of the model of chemical synapses (E_DP_, E_PCN_,I_DP_, I_PCN_, E_CDP_ in Fig 1). The adjustment of the parameters of these synapses is essential to generate the four types of SPI patterns showed by the electromotor command network. An ad hoc iterative tuning of the parameters of the model was performed to match previously described dynamics in the electromotor command network: (i) DP/PCN units firing sporadically before CN, (ii) DP/PCN remaining silent for tens to hundreds of milliseconds after an action potential from CN and (iii) VPd firing burst of action potentials during DP/PCN silence starting *≈*1-8 msec after CN activation.

The intrinsic complexity of manually tuning all the parameters to reflect these dynamics shown by the real network leaded to develop an automatic method for tuning synaptical parameters of the model. This method is described in the next section.

### 2.4 Automatic selection of synapse parameters

An automatic approach was used to tune the parameters of the model synapses. This approach was followed to overcome the lack of specific physiological information about the characteristics of the synapses in the real system. A genetic algorithm (GA) was developed and applied to evolve the parameters of the synapses in order to reproduce the variability of the electromotor command system patterns.

Each individual *I* in the population of the GA was conformed by a set of 20 parameter values: *α, β, g, t*_*max*_ for each of the 5 synapses of the model (see Fig. 1, Eq. 4 and Eq. 5). Each *I* in a generation had different randomly modified values from the initial ad hoc tuned model. Alterations for each parameter were constrained under distinct ranges in different executions of the GA.

Each generation, starting from the initial one, was formed by 100 different individuals. The GA follows an steady state GA scheme [1, 22]. In this scheme, a temporary population was created by cross and mutation and it was added to the original population. All individuals were then evaluated (using the fitness function described in the next section) and ranked according to their grade. The worst individuals were discarded in order to return the population to its original size. The best individuals (10%) were maintained between generations. This process continued for a predefined number of generations or until a certain fitting value was reached.

#### 2.4.1 Fitness function

Each individual, defined by a set of values for the previously specified parameters, was evaluated being simulated under a predefined set of 4 different simulation cases (*S*), each one corresponding to a target SPI: Acceleration (*S*_*acc*_), scallop (*S*_*sca*_), rasp (*S*_*rasp*_), cessation (*S*_*cess*_). Simulation cases (*S*_*pat*_) established the current inputs required to reproduce the *pat* pattern. Each individual *I* was modeled under all four simulation cases. The fitness function of the overall individual (*f* (*I*)) was defined as the sum of the fitness results under each case:

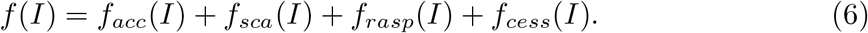

The four target patterns (acceleration, scallop, rasp and cessation) were defined in terms of an ordered sequence of IPIs (*p*_0_, …, *p*_*m*_) where *p*_*i*_ is each IPI in temporary order. The output of a simulation case was also defined in terms of an ordered sequence of IPIs. To compare sequences, they were normalized to the same duration (1000 arbitrary units), regularly interpolated (every 20 arbitrary units, obtaining *n* = 50 points) to detect the pattern shape, and differentiated, as SPI patterns are better defined by the increasing/decreasing slopes between IPIs better than the absolute timing values (see Fig. 3 for an example). The pseudocode of this fitness function is shown in Table 1.

**Table 1.**
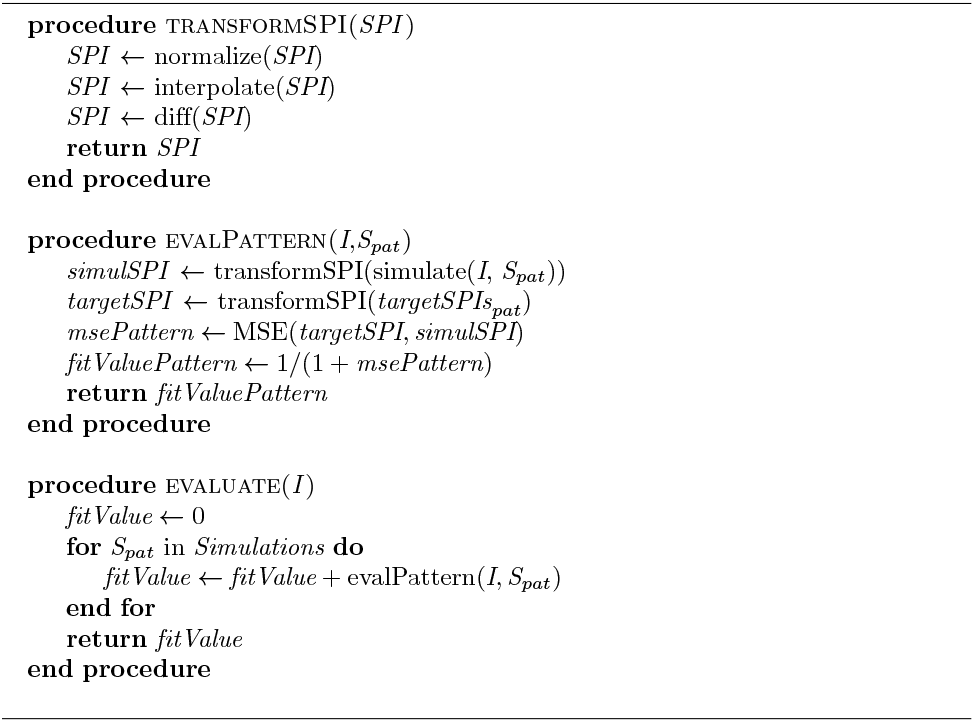
Pseudocode of the fitness function, which relies on three different procedures: First, *tranformSPI* procedure considers the list of sequential IPIs that form one SPI and applies the transformations showed in Fig. 3. Then, *evalPattern* procedure applies *tranformSPI* to both the simulated SPI (*simulSPI* procedure) and the target pattern (*targetSPI*) to compare them by calculating the mean squared error between both (*msePattern*, calculated as described in Eq. 7). The *evalPattern* procedure returns the fitting value of a pattern (*fitValuePattern*) calculated as described in Eq. 8. Finally, *evaluate* procedure invokes *evalPattern* for each of the four patterns (scallop, acceleration, rasp and cessation) and adds them to return the fitness value of the overall individual (*fitValue*, equivalent to *f* (*I*) in Eq. 6).

**Table 2.** Relevant synapse parameters of the GA adjusted to synthetic patterns (S-GA) and GA adjusted to recorded *Gnathonemus Petersii* patterns (R-GA, see Fig. 2) configurations of the model (*threshold* = 0; *E*_*syn*_ = *-*80; T = 1; see Eq. 4 and Eq. 5).

| **S-GA** |  |  |  |  |
| --- | --- | --- | --- | --- |
| Synapse | $\alpha$ | $\beta$ | $g_{syn}$ | $t_{max}$ |
| I<sub>DP</sub> | 9.05623 | 0.00272207 | -0.1251 | 223.097 |
| I<sub>PCN</sub> | 7.50072 | 0.0270961 | -0.27628 | 169.763 |
| E<sub>DP</sub> | 4.949 | 0.127261 | 0.179499 | 78.9001 |
| E<sub>PCN</sub> | 4.949 | 0.148451 | 0.259499 | 78.9001 |
| E<sub>CDP</sub> | 4.79499 | 0.00327418 | 0.705623 | 396.811 |

| **R-GA** |  |  |  |  |
| --- | --- | --- | --- | --- |
| Synapse | $\alpha$ | $\beta$ | $g_{syn}$ | $t_{max}$ |
| I<sub>DP</sub> | 0.537866 | 0.0052969 | -0.165774 | 177.288 |
| I<sub>PCN</sub> | 5.94833 | 0.00129492 | -0.307652 | 167.175 |
| E<sub>DP</sub> | 5.98168 | 0.120037 | 0.238086 | 9.51458 |
| E<sub>PCN</sub> | 5.02672 | 0.218571 | 0.199753 | 84.4537 |
| E<sub>CDP</sub> | 4.43346 | 0.0137051 | 0.647143 | 428.988 |

**Figure 3.**
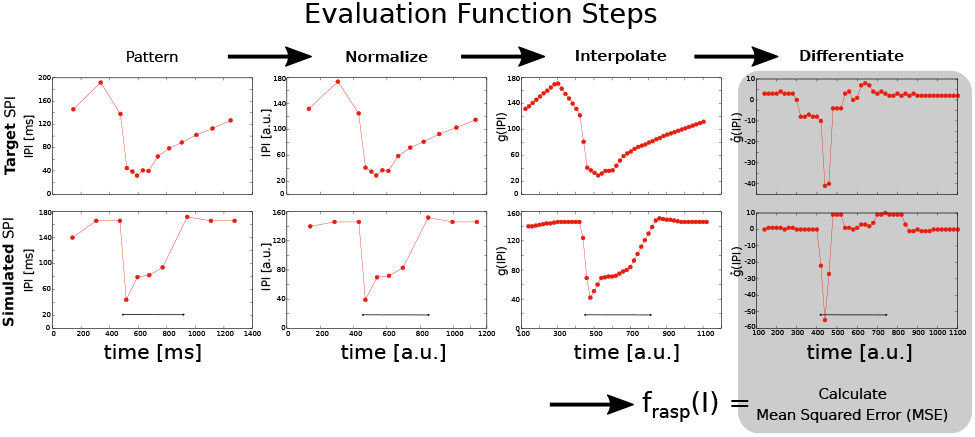
Steps for SPI output fitness evaluation in a rasp pattern example comparing a target SPI (top) and a simulated SPI (bottom). The input current step function is injected at 500 ms and lasts for 400 ms. First, the SPI is normalized to 1000 arbitrary units (in the figure, it is represented starting at the first IPI). Normalized IPIs are then interpolated every 20 ms and differentiated (*ġ*(*IP I*)). Finally, the mean squared error is calculated between the target SPI pattern and the simulated one.

Finally, the fitting value *f*_*pat*_(*I*) was calculated by comparing the target SPI patterns after these transformations 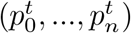 with the SPI model outputs after the same transformations (*p*^*S*^(*I*)_0_, …, *p*^*S*^(*I*)_*n*_) using the mean squared error (*MSE*) as follows:

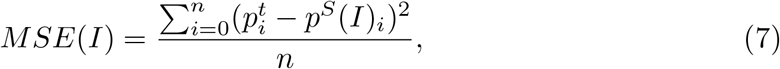

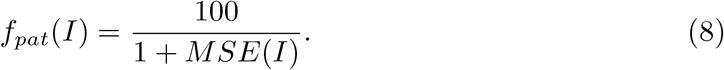

#### 2.4.2 Model simulation

Since the model presented here is multi-objective (i.e., it reproduces different SPI patterns when modifying only the model inputs), a set of four different simulation cases (each one related to a distinct SPI pattern) was determined. Each simulation case was defined by the input current values received by the model (In_VPd_, In_DP_, In_PCN_ in Fig. 1) during the simulation of the four SPI patterns shown in Fig. 4.

**Figure 4.**
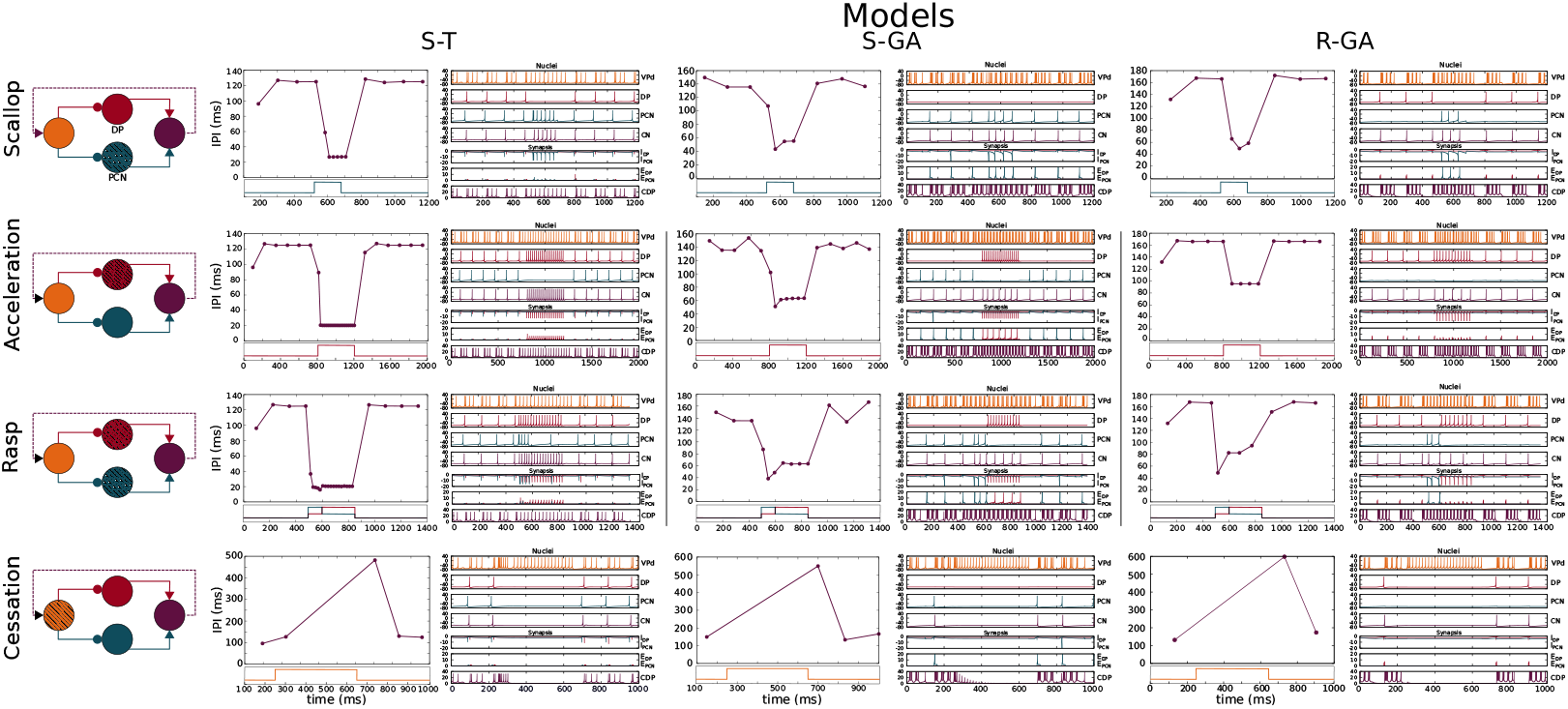
Simulation of the four SPI patterns in three different model configurations: *ad hoc* tuned configuration (S-T), a configuration optimized to synthetic examples (S-GA), and a configuration adjusted to patterns recorded from freely-behaving specimens (R-GA). Each row showed (left) a schematic of the network where the nucleus/nuclei responsible for generating the SPI [6] is/are highlighted using a stripped pattern. The results for each configuration are depicted in two columns: the first one (left) shows SPIs resulting from the simulation and the second one (right) shows the nuclei voltages and synaptic currents. Each chart is related to its corresponding nuclei/synapse by color. Step functions used as current inputs to activate nuclei/nucleus are depicted under each SPI pattern, and they are also related to their corresponding nuclei by color. The model parameters are described in B. Simulation parameters used in the simulations are described in C.

This predefined inputs were step functions that represent projections received by VPD, DP and PCN nuclei from other sources of the nervous system. We assume that with no input, there is no activity in the model network. Before each simulation, the model was initialized for a random amount of time. During this period, the model is stimulated to reproduce a base rhythm of IPIs of*∼*120ms (in the range of IPIs showed by the real fish during resting). Input values for each SPI case are described in C. The source code of both the electromotor command network model and the GA for synaptic parameter optimization are provided (see G). The GA can be easily adapted to any alternative experimental data from weakly electric fish.

### 2.4.3 Robustness to input variability and overfitting analysis

An analysis for all the different configurations of the model was conducted to assess robustness to input variability and discard overfitting to the predefined stimulation cases. The step functions used as inputs in the simulations are meant to reproduce the nuclei activation associated with each SPI (see C). As a consequence, similar SPIs were expected to result from distinct inputs as long as the appropriate nucleus was stimulated. According to this hypothesis, a robustness analysis to input variability was conducted.

To perform this analysis, a set of different simulations was conformed by modifying the intensity and duration of the predefined model inputs (i.e., the inputs used in the GA for simulating the patterns showed in Fig. 4) up to a 50%. Being 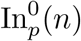(*n*) the predefined value of the input received by nucleus *n* in the *p* pattern simulation case (see C), and 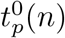(*n*) the duration of the step, the set of simulations to assess robustness is built by modifying intensity from 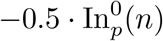(*n*) to 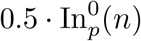(*n*) in steps of 0.05, and also modifying duration from 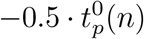 to 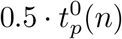 using the same interval steps (see Fig.5 - Right). Then, the reproducibility of the patterns under this set of stimulation cases was evaluated using the fitting function. In this case, a relative fitting value was defined as the change ratio of the fitting value from the default stimulation case. Being *f* ^0^ the fitness value of the model under predefined stimulation, and *f*^*i*^ the fitness value under a given stimulation case of the set, then, the relative fitting value (Δ*f* (*i*)) was given by:

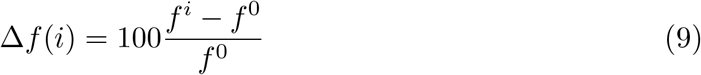

where negative results mean a decrement in the fitting value when varying the stimulation inputs. When *f*^*i*^ = *f* ^0^, Δ*f* (*i*) = 0, which is the case in the central point of robustness charts that we will discuss later (see Fig. 5 - Left and Fig. 6)Robustness of the model is evaluated below in terms of the relative increments and decrements of this value. Strong robustness is defined as the ability to maintainΔ*f* (*i*) *>* 0. Note that, here, the fitness function was used again to measure the distance to target patterns, but no GA was employed.

**Figure 5.**
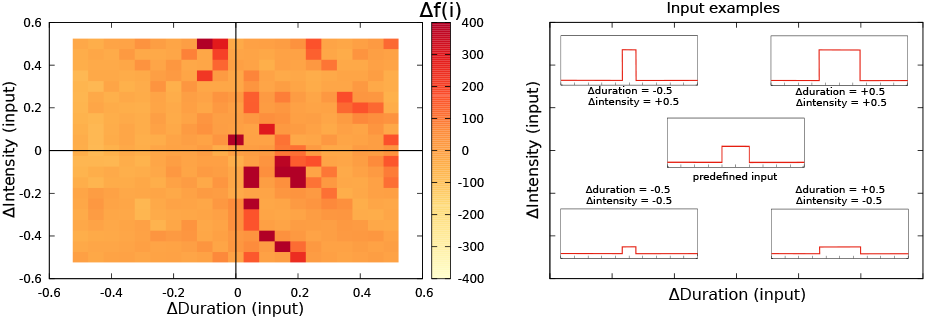
Left: Robustness analysis to input variability of the R-GA electromotor model tuned to recorded Gnathonemus Petersii SPI patterns. The central point in the left panel represents the reference fitness value, the one obtained simulating the model under the predetermined simulation conditions (i.e., ΔDuration = 0 and ΔIntensity = 0). Duration and intensity of the step function current input was variated from −0.5 to 0.5 from their initial values and the relative change in the fitness value was calculated as described by Eq. 9. White and lighter yellowish colors represent decreases in the fitness value under variable stimulation. Orange colors represent an equivalent result to the reference fitness value. Darker red colors represent an improvement in the fitness value. Right: Representative example of distinct current inputs in the simulation at different places of the chart.

**Figure 6.**
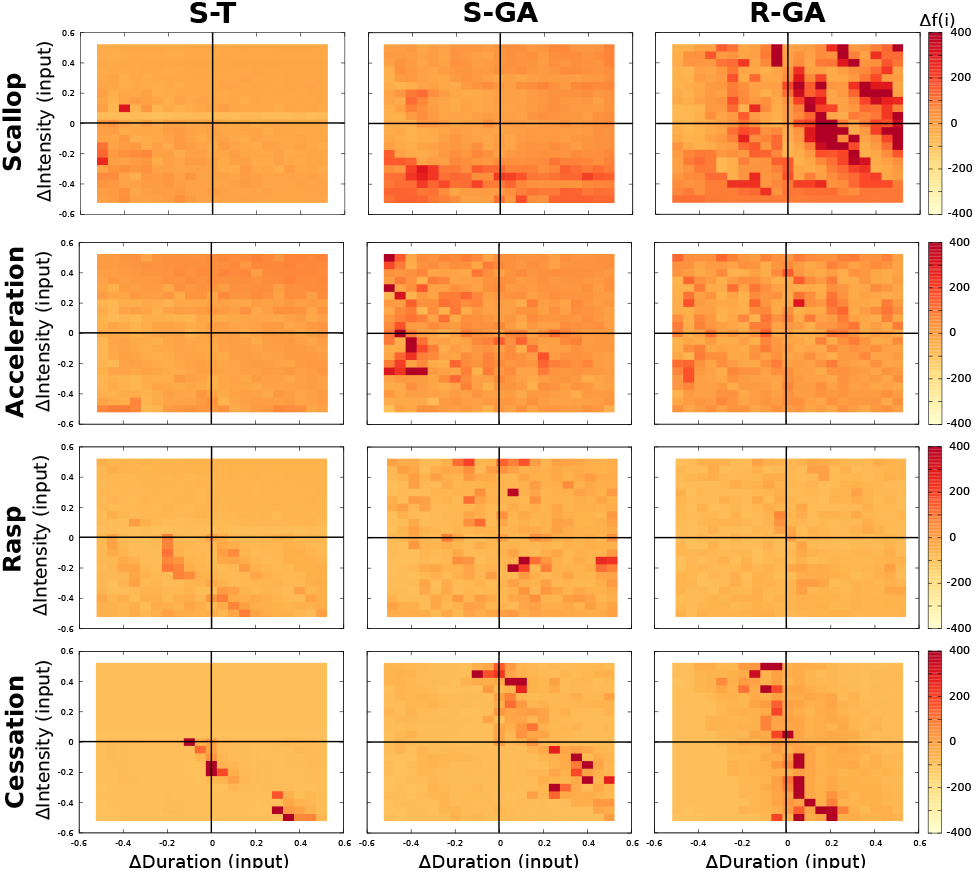
Deviation from the 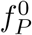 (reference fitness value) of each model configuration (ST, S-GA and R-GA) modifying default current input intensity 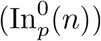 from 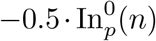 to 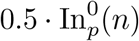 in steps of 0.05, and also modifying default current input duration 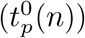 from 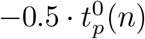 to 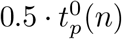 using the same step. The color scheme is the same as the one described in Fig 3. See Fig. 7 for a representation of the IPIs (mean and variance) resulting from worse (light yellow), similar (orange) and better (dark red) fitness results in R-GA.

## 3 Results

Three different configurations of the model were analyzed (see Table 2 and also B), first the ad hoc tuned configuration (S-T) and the other two from GA-tuning of synaptic parameters, one fitted to synthetic SPI patterns (S-GA) and the other one fitted to reproduce SPI patterns recorded from *Gnathonemus Petersii* specimens (R-GA) (See Fig. 2). Each configuration reproduced all four SPI patterns with a different level of accuracy and robustness. A specific SPI pattern was evoked only due to changes in the inputs (In_*VPd*_,In_*DP*_,In_*PCN*_ in Fig. 1) without modifying any internal parameters. GA configurations showed some shared characteristics between them when compared with the S-T configuration, as the increases in the absolute value of the synaptic conductances (*g*_*syn*_). Note that some of these parameters noticeable change, this is the case of the *g*_*syn*_ parameter for the *E*_*P CN*_ synapsis (an increase of four times for S-GA and five times R-GA from its initial value in S-T, as it can be seen in Table 2 and B). These similarities are remarkable as network parameters are fitted to a different set of samples of the same SPI patterns. Also remarkable is that S-GA showed similar *α, β* and *t*_*max*_ parameters for both *E*_*DP*_ and *E*_*P CN*_ projections (Fig. B), which means that the timing of the synapses is almost symmetrical for both excitatory pathways (Fig. 1).

Four different SPIs patterns were simulated (Fig. 4) using the three configurations of the model: S-T, S-GA and R-GA.

Scallop simulations in both S-GA and R-GA showed the typical behavior associated to this pattern (which is a sudden drop to short IPIs (around 40 ms) followed by an almost immediate recovery). The shorter IPI reached in S-T was slightly faster (*∼*25 ms), although the burst duration in all the three configurations remained almost the same. Both in S-GA and R-GA, scallop SPIs reached lower IPI duration than accelerations, in accordance with the activity recorded from the fish.

In all the three configurations, accelerations (Fig. 4 − B) showed a series of almost regular shorter IPIs. In S-T, short IPIs during acceleration were around 20 ms. In S-GA, IPIs during acceleration were longer (around 60 ms. in S-GA and 85 ms in R-GA), better complying with the accelerations from the fish (which are highly variable, but consistently larger than 20 ms, and even larger in *Gnathonemus Petersii* recordings). It is worth noting that, contrary to what happened in the initial S-T model, in both GA-fitted configurations CN integrated several DP spikes before firing. Regarding IPIs regularity during the acceleration, IPIs within SPI had approximately the same duration. Both R-GA and S-GA configurations showed regularity in the sequence of short IPIs that define acceleration, although in S-GA the starting IPI tended to be slightly shorter.

Regarding S-T results, scallop and rasp SPI patterns cannot be easily distinguished, as both showed acceleration-like short regular IPIs (Fig. 4). The total SPI duration was different for each pattern, but in S-T rasps and scallops lacked their own characteristic internal structure.

In S-GA and R-GA, rasps (Fig. 4 - C) showed an initial scallop-type decrease to IPIs around 40 ms, followed by a sustained burst of regular short IPIs like in accelerations. As mentioned before, in S-T the pattern was not fully identifiable. In S-GA the IPIs were shorter, so the SPI was formed by a larger number of IPIs. Also, the recovery to larger IPIs was more abrupt in this case. Again, in both GA-fitted configurations, CN integrated several DP spikes in their IPIs. Conversely, PCN spiking was tightly phase-locked with CN.

Finally, cessations (Fig. 4 - D) showed the expected stop in the generation of pulses during long time periods (of around 500 ms) in all three configurations.

Results for the robustness analysis to input variability of R-GA are depicted in Fig. 5 using a color representation of Δ*f* (*i*) as described by Eq. 9. Results did not show relevant decreases in the fitting value for changes up to 50% in the intensity and duration of the default simulation values. Conversely, they showed a limited increase of the fitting value when slight decreases in duration were balanced with slight increases of intensity and, similarly, when slight increases in duration were balanced with slight decreases of intensity.

To discard overfitting to any of the SPI patterns, results of this analysis disaggregated by pattern are also shown in Fig. 6 (rightmost column). No relevant drops in the fitness results stands out in these panels for variations of up to 50% in the intensity and duration of the predefined stimulation inputs. Quite the opposite, better evaluation results (depicted in dark red) were usually obtained, for example, in R-GA scallops, specially for increases in the input duration. For comparison, we also depict in Fig. 6 the robustness results of the other two configurations of the model (those adjusted to synthetic patterns: S-T, S-GA). S-T moderately improved the fitness results in accelerations, and also in cessations when a increase in the duration of the inputs were compensated with a decrease in the input duration. S-GA, in general, showed greater robustness to input variability, maintaining or improving fitness in all SPI patterns.

In Fig. 7, IPI mean and standard deviation of each SPI pattern simulated during the robustness analysis of the R-GA model are depicted. Line depicted in red is the mean SPI of the several executions with variable inputs and the red shade represents the standard deviation of the results. Finally, black line corresponds to the closer target pattern used to calculate the fitness. The simulation of each SPI pattern were divided in three partitions according to its fitting value: Worse fit than the one obtained with default inputs (Δ*f < −*100); similar fit (*-*100 *<* Δ*f <* 100) and worse fit (Δ*f >* 100).

**Figure 7.**
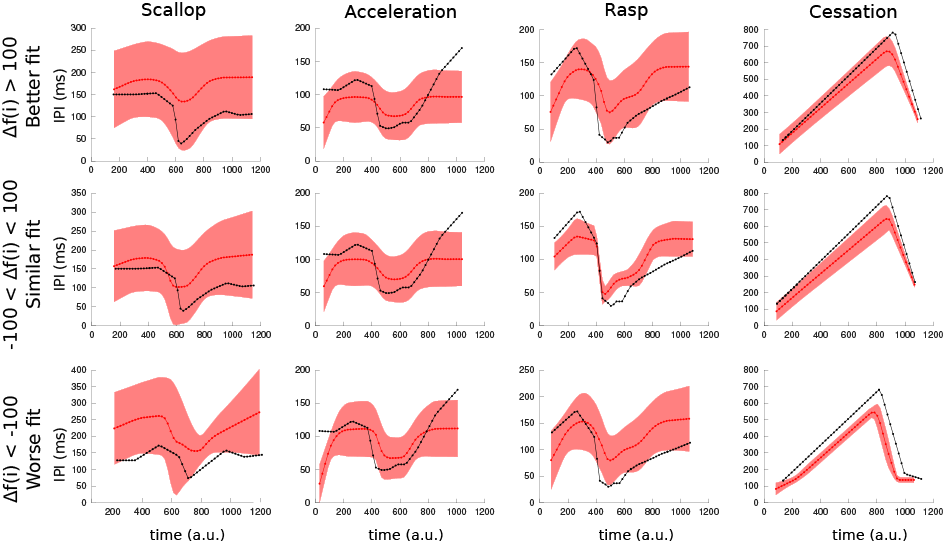
Mean and variance of simulated SPIs using the R-GA model under the variated simulation conditions of the robustness analysis (see Fig. 6). As in the GA fitness function, the duration of each simulation was normalized to 1000 arbitrary units for representation, starting at its first IPI. Red line is the mean SPI from the executions with variated inputs, red shade represents the standard deviation. Finally, the SPI represented in black is the closer target pattern. Simulations of each SPI pattern were divided in three partitions according to its fitting value: the ones that obtains better fitting (upper row: Δ*f <-*100), those that obtains similar results to the reference fitness value (center row: *-*100 *<* Δ*f <* 100) and the ones that yield worse fitting results (bottom row: 100 *<* Δ*f*).

## 4 Discussion and conclusions

Computational models have been previously used to answer different questions in the study of electroreception and electrogenesis. Regarding electrogenesis, authors in [24, 27, 31] used an anatomically detailed model of the pacemaker of *Apteronotus leptorhynchus* to study the electric organ signal and its spatiotemporal features in wave-type fish. In [30] a model for the *Eigenmannia* was employed to address the jamming avoidance response in these fish. Electroreceptor models are of particular interest as they have a relevant role in understanding the electrical sense, even in non-electroactive species. On the other hand, bioinspired approaches intend to mimic electrolocation by building robotic electrosensory systems at different abstraction levels. Aiming to develop an underwater autonomous robot with electric sense, a robotic model of the electric fish fins was developed to study fish manoeuvrability [28].

Despite the variety of models regarding signal generation in weakly electric fish, the development of a computational model of the electromotor command network that can produce the variability of SPIs observed in pulse mormyrids had not been attempted until now. The model presented here assesses the relevance of different parameters in the electromotor command network for reproducing as a whole the diverse temporal structure of output patterns displayed by the living system. Our results suggest that the diversity of SPIs shown by the system is only possible due to a dynamic balance of intensity and timing between the synapses of the network triggered by input stimulation without temporal structure. This highlighted the hypothesis that relevant synaptic parameters (and not only the nucleus dynamics or the network topology) play an important role for reproducing the whole set of SPIs observed in experimental data.

A multi-objective genetic algorithm (GA) was developed and applied to tune the synaptic parameters of the model. GA optimization method allowed us to readily obtain different parameter configurations which optimize the ability of the model to reproduce all target patterns. Attending to the results, automatic synaptic parameter optimization allowed to prove that it is possible to reproduce the four preselected SPI patterns using the described topology.

Robustness results show that overfiting to the stimulation inputs was avoided. Although there are considerable drops in fitness for certain combinations of inputs, the general tendency was to maintain (and sometimes even improve) the performance of the model under distinct stimulation (as shown in Fig. 5 and Fig. 6). S-GA and S-T configurations showed a strong robustness to changes in the stimulation inputs (both in terms of intensity and duration), which is coherent with the fact that synthetic target patterns showed less richness and thus were more easily reproduced. In R-GA, rasps were the least robust SPI patterns, but even those with the worsts fitting results kept a recognizable, though slightly flattened, rasp shape. These robustness results give credit to the idea that SPI generation depends on which nucleus is activated and not so much on the intensity or the duration of the inputs.

Regarding the resulting model configurations (described in B), they all indicate that the corollary discharge must have a larger maximum release time than any other of the synapses in the model. This characteristic is always present in all best individuals obtained from the GA and is coherent with the idea that it might be an indirect pathway. Also, the fact that S-GA showed similar *α, β* and *t*_*max*_ parameters for both *E*_*DP*_ and *E*_*P CN*_ projections (see table 2) reinforces the hypothesis of synaptic intensity being the decisive parameter in the elicitation of accelerations or scallops. Nevertheless, the conductance ratio between these synapses is not the same in R-GA. It denotes that the relation between synaptic strengths determines the output of the system in a non-linear manner.

Its important to note that SPI patterns are generated over a background of ongoing activity in the electromotor system. This activity is highly affected by noisy external inputs. Although the resting rhythm of the system (IPIs ranging from *∼*100-300 msec., following a bimodal distribution) is easily reproduced by the model using biologically plausible input values, an in-depth study of the baseline activity contribution to the overall output (and how it affects SPI temporal structure) is yet to be done.

The implemented model and its computational efficiency also enables closer-to-natural stimulation to perform more realistic closed-loop experiments with the real system (as pulse-type mormyrids have been previously used in several closed-loop studies where stimulation is guided by the fish own activity, like in [14, 15, 21]). In this context, we can expect these experiments to benefit from a more realistic stimulator based in the described model [21]. The model can also allow further studies of the underlying mechanisms of electrocommunication, although the internal relationship between fish skin electroreceptors and the electromotor system has yet to be further studied.

All the software developed and used in the analysis presented here (the model, the multi-objective GA used for parameter adjustment with the fitness function used to represent and compare different sequences of temporal firing patterns and the robustness analysis) is provided for future studies of this network and similar networks of other animal models (see S7 supplementary material). These tools can deepen our understanding about the role of the ensemble connectivity and the synaptic mechanisms shaping functional neural temporal structures.

## Acknowledgments

This work was supported AEI/FEDER grants TIN2017-84452-R, PGC2018-095895-B-I00. The funders had no role in study design, data collection and analysis, decision to publish, or preparation of the manuscript.

## A Target SPI patterns: synthetic and recorded

Synthetic SPI data was used for S-GA tuning to reflect the characteristic temporal structure of reported behavioral SPI patterns [6, 11, 26] as they are described in the section *Characteristic sequences of pulse interval (SPIs)*. Four different synthetic SPIs were generated for each behavioral pattern (scallop, acceleration, rasp and cessation). Figure 2 (bottom) shows only one set of the patterns.

Non-synthetic SPIs were recorded from three different freely-behaving *Gnathonemus Petersii* specimens in a 80 l water tank, using four differential dipoles placed in the tank forming 45° angles between them. The grounding electrode was also located in the aquarium. The signal from the dipoles was amplified (TL082 JFET-Input Dual Operational Amplifier), summed (LM741 Operational Amplifiers), squared (using AD633 Analog Multiplier) and then digitized at 15 Khz by a DAQ board (NI PCI-6251, National Instruments Corporation).

IPIs detection was performed using an spike detection algorithm as described in [21]. Examples of SPI recorded from *Gnathonemus Petersii* specimens are depicted in Fig 2 (top). We selected from the *Gnathonemus Petersii* recordings those SPIs that most closely resemble the temporal structure of SPI patterns reported in other mormyrid species. The evolutionary approach used in this paper can be applied to reproduce different sets of temporal SPIs given an adequate network topology. Permission of the ethics committee of Universidad Autónoma de Madrid was obtained (TIN2017-84452-R/CEI-88-1661). All experiments were noninvasive behavioral trials. All animals behaved normally after the experiments.

## B Model parameters

The electromotor system was modeled using four nuclei and five synapses. The nuclei VPd, DP, PCN and CN were simulated using the neuron model developed by Izhikevich (2003) [19], where a wide range of neuron dynamics can be evoked by selecting different values of the parameters *a, b, c* and *d* [13]. These values were tuned ad hoc to reproduce dynamics described by previous neurophysiological studies. The following firing regimes were used:

- Low-threshold spiking (LTS): A type of low firing threshold activity tipically observed inhibitory cortical cells. In this mode, a neuron can fire high-frequency trains of spikes (burst) and it also shows frequency adaptation while the stimulation is sustained (see table 2). The parameters of VPd in the model correspond to this kind of behavior.
- Regular frequency spiking (RFS): This mode correspond to spike firing at a regular frequency when stimulation amplitude is held constant (see table 2). It is the firing regime in which DP operates.
- Regular Spiking (RS): This mode increases its regular firing frequency when stimulated (see table 2). Adaptation to the stimulus results in a recovery from short interspike intervals to the regular firing frequency. PCN follows this behavior.
- Phasic Spiking (PS): This firing regime operates as an integrate-and-fire model, although it fires only a single spike as a response to the input, and remain quiescent afterwards. In the model, it is the firing regime used for CN.

Table 1 show the parameter values used in the electromotor model. Figures in table B.2 show the activity of the individual dynamics of all nucleus types considered in the model.

The synapses (*I*_*DP*_, *I*_*P CN*_, *E*_*DP*_, *E*_*P CN*_ and *E*_*CDP*_) were simulated using the model described by [13]. The most relevant parameters of the model (*α, β*, gsyn and *t*_*max*_) were automatically adjusted to the target SPI patterns using the genetic algorithm (GA) described in this paper. Table 2 showed the initial ad hoc tuned configuration (S-T).

**B.1 Table.** **Neuron parameters** of the model to reproduce the dynamics of the electromotor nuclei as described by previous neurophysiological studies [6] (see Eq. 1,Eq. 2 and Eq. 3).

| Neuron | Dynamics | a | b | c | d |
| --- | --- | --- | --- | --- | --- |
| VPd | Low-threshold spiking (LTS) | 0.02 | 0.25 | -65 | 2 |
| DP | Regular frequency spiking (RFS) | 0.1 | 0.26 | -65 | 2 |
| PCN | Regular spiking (RS) | 0.02 | 0.2 | -65 | 8 |
| CN | Phasic spiking (PS) | 0.02 | 0.25 | -65 | 6 |

**B.2 Table.**
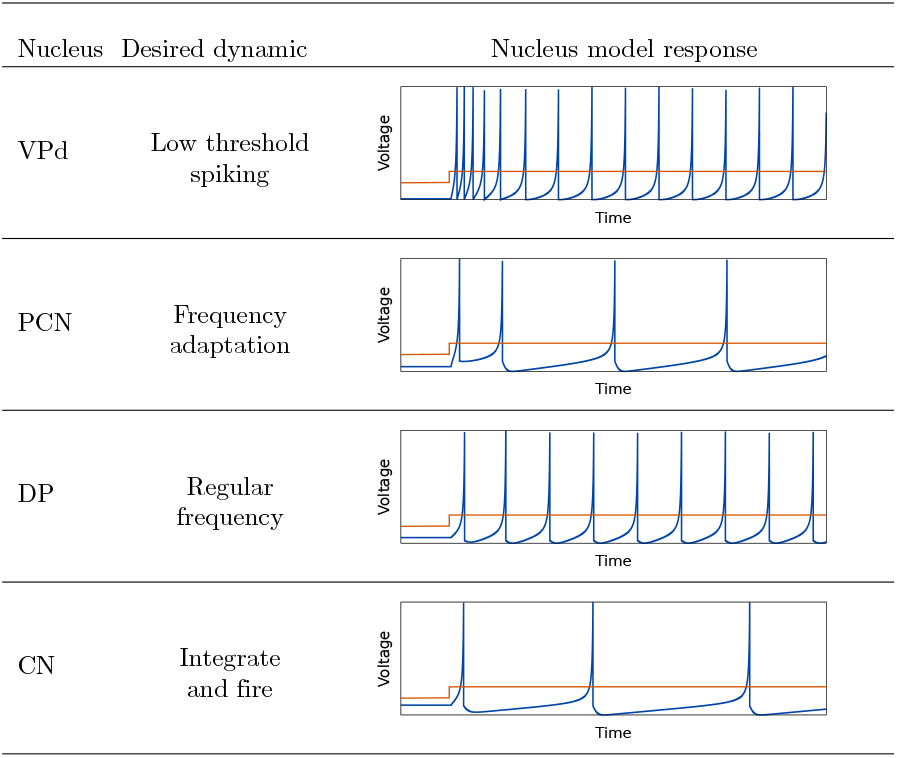
Dynamic properties of the model nuclei chosen in this study to match their biological counterparts. Values for the parameters of the model by [19] were tuned to replicate the desired experimental dynamic (parameter values are detailed in B). Last column shows the activity of each isolated nucleus model in response to an input current step (*I*_*syn*_(*t*) in Eq. 1).

| Nucleus | Desired dynamic | Nucleus model response |
| --- | --- | --- |
| VPd | Low threshold spiking |  |
| PCN | Frequency adaptation |  |
| DP | Regular frequency |  |
| CN | Integrate and fire |  |

**B.3 Table.** Relevant synapse parameters of the ad hoc tuning to synthetic patterns (S-T) configuration of the model (*threshold* = 0; *E*_*syn*_ = *-*80; T = 1; see Eq. 4 and Eq.5).

| Synapse | S-T |  |  |  |
| --- | --- | --- | --- | --- |
| | $\alpha$ | $\beta$ | $g_{syn}$ | $t_{max}$ |
| I<sub>DP</sub> | 5 | 0.005 | -0.12 | 160 |
| I<sub>PCN</sub> | 5 | 0.005 | -0.15 | 160 |
| E<sub>DP</sub> | 5 | 0.1 | 0.1 | 30 |
| E<sub>PCN</sub> | 5 | 0.18 | 0.05 | 30 |
| E<sub>CDP</sub> | 5 | 0.02 | 0.3 | 400 |

## C Simulation parameters

To simulate the electromotor command system, VPd, DP and PCN received external currents, which represent inputs. These predefined values represent current inputs received by the nuclei from other sources of the nervous system. The following tables show the current input values for simulating scallops (C.4 Table), accelerations (C.5 Table), rasps (C.6 Table) and cessations (C.7 Table). The first row in each table represents simulation time (in milliseconds), and subsequent rows represents neuron current input for each nucleus model (VPd, DP, PCN) during this time. These inputs were step functions used to simulate the SPIs shown in Fig. 4. Also, these are the predefined simulation parameters for the GA evolution and the 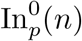robustness analysis (center values in Fig. 5 and Fig. 6).

**C.4 Table.** Scallop simulation parameters. This SPI pattern is evoked by increasing current input to PCN neuron. VPd input (−0.5 mA) and DP input (1.7 mA) are constant in this pattern.

| Time (ms) | [0, 520] | [520, 680] | [680, 1200] |
| --- | --- | --- | --- |
| PCN input | 6.5 | 14 | 6.5 |

**C.5 Table.** Acceleration simulation parameters. This SPI pattern is evoked by increasing current input to DP neuron. VPd input (−0.5 mA) and PCN input (6.5 mA) are constant in this pattern.

| Time (ms) | [0, 800] | [800, 1200] | [1200, 2000] |
| --- | --- | --- | --- |
| DP input | 1.7 | 4 | 1.7 |

**C.6 Table.** Rasp simulation parameters. This SPI pattern is evoked by increasing current input to both DP and PCN neurons. VPd input (−0.5 mA) is constant in this pattern.

| Time (ms) | [0, 500] | [500, 580] | [580, 830] | [830, 1330] |
| --- | --- | --- | --- | --- |
| DP input | 1.7 | 4.5 | 4 | 1.7 |
| PCN input | 6.5 | 15 | 7 | 6.5 |

**C.7 Table.** Cessation simulation parameters. This SPI pattern is evoked by increasing current input to VPd neuron. DP input (1.7 mA) and PCN input (6.5 mA) are constant in this pattern.

| Time (ms) | [0, 250] | [250, 650] | [650, 1000] |
| --- | --- | --- | --- |
| VPd input | -0.5 | 8 | -0.5 |

## D Homogeneous phasic spiking neurons

In order to address the contribution of the ad-hoc tuned nuclei parameters to the overall network output, all nuclei (VPd, DP, PCN, CN) were adjusted to homogeneous parameters reproducing a phasic spiking behavior. GA was then applied to fit synaptical parameters to both recorded and synthetic data. Fitting results (calculated using the evaluation function) were consistently lower than those obtained in S-GA and R-GA. The best result obtained is depicted in Fig. 1. In this case all SPI patterns were still recognizable. This denotes that SPIs temporal structure is not directly determined by the nuclei intrinsic characteristics. On the contrary, this reinforces the idea that it is synaptic conductances in the mormyrids electromotor system which play a primary role in the generation of the temporal structure of SPI patterns.

**Figure 1.**
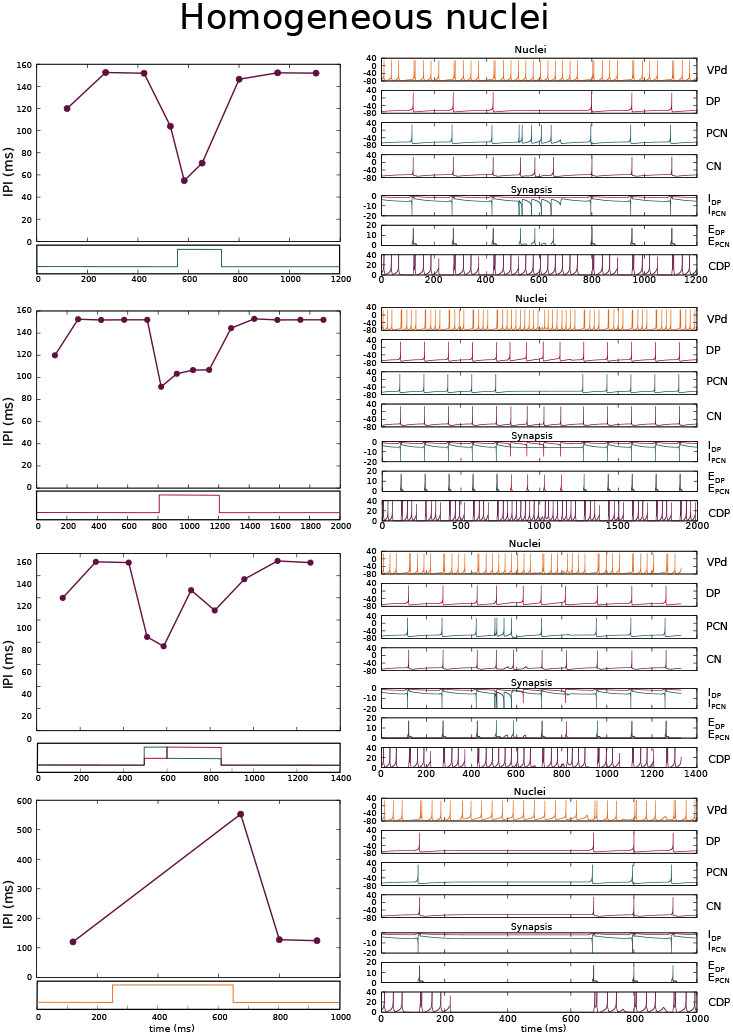
Simulation of the four SPI patterns in the best fitting model using homogeneous nuclei parameters reproducing a phasic spiking behavior. The results are depicted in two columns: the first one (left) shows SPIs resulting from the simulations and the second one (right) shows the nuclei voltages (top) and synaptic currents (bottom).

## E No corollary discharge

In order to address how the presence of a corollary discharge pathway (CDP) contributes to the variability of SPI patterns showed by the mormyrids electromotor system, CDP synapse was removed from the original model. GA was then applied to fit synaptical parameters (for the rest of synapses) to both recorded and synthetic data. Fitting results calculated using the evaluation function were consistently lower than those obtained in S-GA and R-GA. The best result obtained in our tests is depicted in Fig. 1, where SPI patterns are not recognizable anymore. It is relevant to note that cessation SPI pattern becomes not reproducible, even though the same current input was applied to VPd (which provides inhibition to DP-PCN).

**Figure 1.**
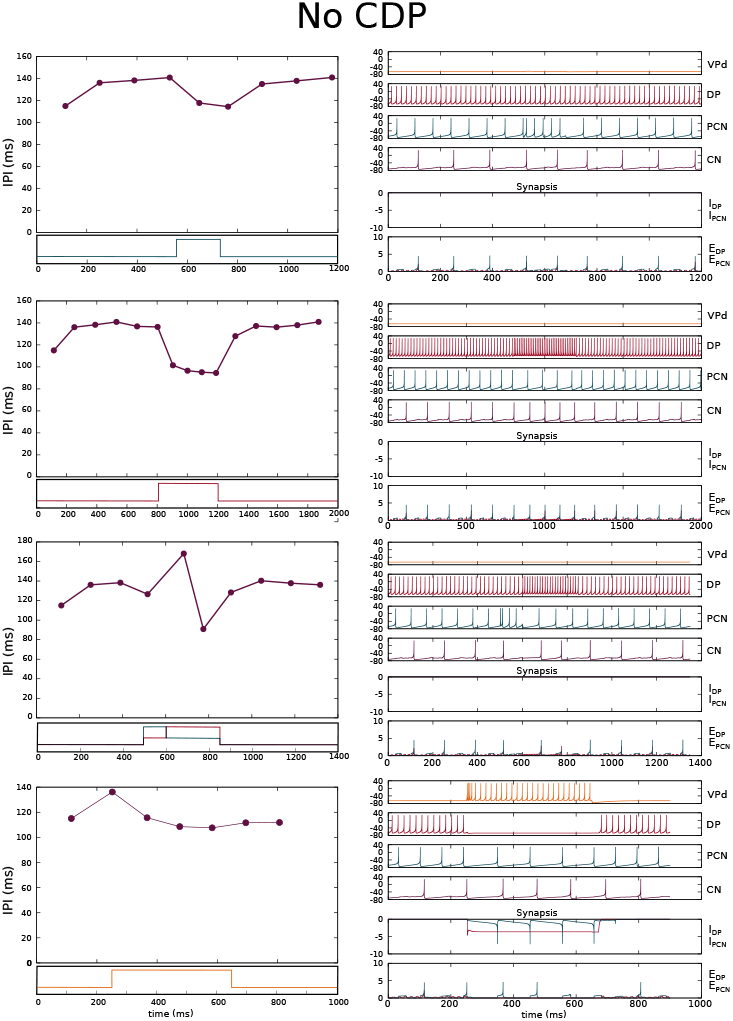
Simulation of the four SPI patterns in the best fitting model without corollary discharge pathway. The results are depicted in two columns: the first one (left) shows SPIs resulting from the simulations and the second one (right) shows the nuclei voltages (top) and synaptic currents (bottom).

## F No PCN pathway

In order to address how the topology of the model and, more especifically, the presence of two different synaptical pathways affects the diversity of SPI patterns showed by the mormyrids electromotor system, one of these two pathways was removed (more precisely, the one corresponding to PCN nucleus). GA was then applied to fit synaptical parameters (for the rest of synapses) to both recorded and synthetic data. The best result obtained in our tests is depicted in Fig. 1, where SPI patterns are not recognizable anymore. This reinforces the importance of the two synaptical pathways in the topology of the model to reproduce the variability of SPIs showed by the electromotor command network.

**Figure 1.**
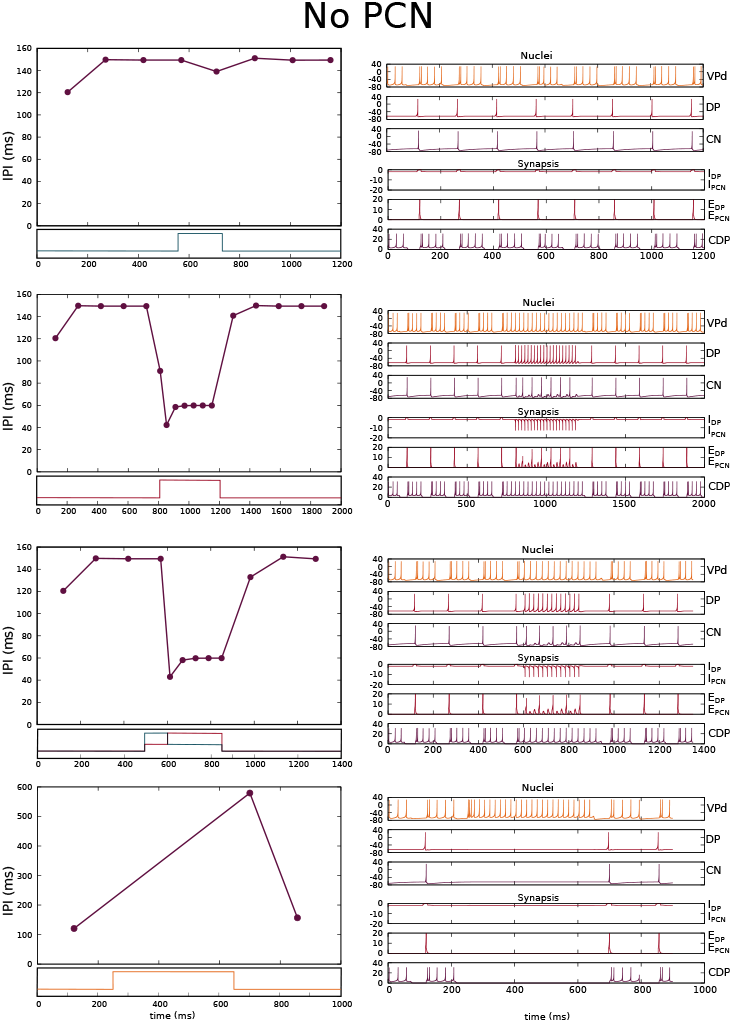
Simulation of the four SPI patterns in the best fitting model without PCN nucleus pathway. The results are depicted in two columns: the first one (left) shows SPIs resulting from the simulations and the second one (right) shows the nuclei voltages (top) and synaptic currents (bottom).

## G Source code

All the software used to develop and run the model, as well as the model itself, are publicly available under an open-source license in a github repository^1^.

The repository includes C++ code needed to run the model, which makes use of Neun dynamical-systems library^2^. The genetic algorithm applied to automatically tune the parameters of the model, implemented using GAlib^3^, it is also provided. Some examples of the recorded and synthetic patterns used as targets to tune the model are also included. Finally, the repository also includes detailed instructions to compile and use the software.

## Footnotes

1 https://github.com/GNB-UAM/electromotor-nmodel

3 https://code.launchpad.net/∼elferdo/neun/trunk

3 http://lancet.mit.edu/ga/

## Notes

https://github.com/GNB-UAM/electromotor-nmodel

### Competing Interest Statement

The authors have declared no competing interest.

### Summary of Updates

The following appendices has been added to the manuscript: -Homogeneous phasic spiking neurons -No corollary discharge pathway -No PCN pathway Discussion and conclusions has been correspondingly updated.

